# Analysis of metagenome-assembled genomes from the mouse gut microbiota reveals distinctive strain-level characteristics

**DOI:** 10.1101/2020.01.29.926196

**Authors:** Shenghui Li, Siyi Zhang, Bo Li, Shanshan Sha, Jian Kang, Peng Li, Aiqin Zhang, Qianru Ji, Qingbo Lv, Xiao-Xuan Zhang, Hongbo Ni, Xiuyan Han, Miao Xu, Guangyang Wang, Wenzhe Zhang, Yuanyuan Sun, Roujia Xu, Yi Xin, Qiulong Yan, Yufang Ma

## Abstract

The laboratorial mouse harbors a unique gut microbiota with potential value for human microbiota-associated studies. Mouse gut microbiota has been explored at the genus and species levels, but features rarely been showed at the strain level. The identification of 833,051 and 658,438 nonredundant genes of faeces and gut content samples from the laboratorial C57/BL mice showed over half of these genes were newly found compared to the previous mouse gut microbial gene catalogue. Metagenome-assembled genomes (MAGs) was used to reconstruct 46 nonredundant MAGs belonging to uncultured specieses. These MAGs included members across all phyla in mouse gut (i.e. Firmicutes, Bacteroidetes, Proteobacteria, Deferribacteres, Verrucomicrobia, and Tenericutes) and allowed a strain-level delineating of the mouse gut microbiota. Comparison of MAGs with human gut colonies revealed distinctive genomic and functional characteristics of mouse’s Bacteroidetes and Firmicutes strains. Genomic characteristics of rare phyla in mouse gut microbiota were demonstrated by MAG approach, including strains of *Mucispirillum schaedleri, Parasutterella excrementihominis, Helicobacter typhlonius*, and *Akkermansia muciniphila*.

**Importance:** The identification of nonredundant genes suggested the existence of unknown microbes in the mouse gut samples. The metagenome-assembled genomes (MAGs) instantiated the specificity of mouse gut species and revealed an intestinal microbial correlation between mouse and human. The cultivation of faeces and gut contents sample validated the existence of MAGs and estimate their accuracy. Full-length 16S ribosomal RNA gene sequencing enabled taxonomic characterization. This study highlighted a unique ecosystem in the gut of laboratorial mice that obviously differed with the human gut flora at the strain level. The outcomes may be beneficial to researches based on laboratorial mouse models.

## Introduction

The gut microbiota is a dense and diverse ecosystem (1). The associations between altered gut microbial composition and various pathogenesis, such as obesity (2, 3), type 2 diabetes (4), rheumatoid arthritis (5), and allergy (6), has become a research hotspot. Murine models, especially mouse, are widely used in biomedical study. Many studies have showed that mouse intestinal models also play a pivotal role in human gut microbial research. Some phyla in mouse intestinal flora, such as Firmicutes, Bacteroidetes, Proteobacteria, has resemblances with that in human (7, 8). In addition, there are plenty of similarities in anatomy, genetics and physiology between mouse and human (9). These features make the mouse one of the most essential model animals in laboratory. However, mouse gut microbiota could be influenced by many factors, including diet, ambient temperature and cleanliness. Hence, there are still some distinctions between human and mouse gut microbiota. Previous studies have showed that over 50% of genera in mouse gut microbiota are not found in human gut (10, 11). Some human gut-dominant genera, including *Faecalibacterium*, *Prevotella*, and *Ruminococcus*, rarely occurred in the mouse gut microbiota, whereas the mouse gut-dominant genera, such as *Lactobacillus*, *Turicibacter*, and *Alistipes*, were underrepresented in human intestinal flora (9).

Although there are some recent researches concerning bacterial mutations in laboratory mouse (12), few studies had focused on strain-level characteristics of laboratory mouse gut microbiota. In this study, using a metagenome-assembled genome (MAG) approach, we analyzed the microbial genomes that were obtained from laboratorial mouse gut microbiota, and revealed distinctive genomic and functional features of these genomes at the strain level. Our findings suggested a conceivable unique ecosystem in laboratorial mice gut and might benefit the research fields with applications to such model organisms.

## Results

### Microbial contents in the mouse intestinal tract

To investigate the overall gut microbial composition of laboratorial mouse, we collected the faeces and gut content samples from three male C57/BL mice and generated a total of 33.9 Gbp high-quality data (5.7±0.5 Gbp per sample, **Table S1**) via whole-metagenome shotgun sequencing. *De novo* assembly and gene prediction of these metagenomic data generated two nonredundant protein-coding gene catalogues for the faeces and gut content samples, containing 833,051 (average length, 695bp) and 658,438 (average length, 708bp) genes, respectively. Over 50% of genes were shared between two body sites (**Figure 1A**), in agreement with the previous studies showing an enormous commonality of microbial content of different intestinal tract sites (13). When combined with the most comprehensive nonredundant catalogue (representing ~2.4 million genes) established by 184 mice faecal samples from overall the world (14), the current mouse gut microbial gene catalogue therefore contained ~2.8 million genes (**Figure 1A**). A high proportion of new genes (52.2%) in this study might be due to more data amounts per sample, improved assembly methods, as well as the population-specific signatures of the mouse intestinal microbiota (14, 15). This combined gene catalogue allowed an average of 73% reads mapping rate for current sequenced samples, and the remaining reads were generally derived from non-coding zone or unassembled sequences. Notably, based on taxonomic annotation via the NCBI-nt database, 63.6% genes in the mouse gut catalogue could be classified at the phylum level, however, only 8.4% genes could be assigned into a genus and only 6.0% genes could be assigned into a species (**Figure 1B**). At the phylum level, the dominant phylum in mouse intestinal samples were Bacteroidetes (average relative abundance = 28.5%), Firmicutes (23.2%), Proteobacteria (18.4%), Deferribacteres (12.3%) and Verrucomicrobia (4.7%), while at the phylum-level unclassified genes consisted an average of 12.3% abundance in all samples. At the genus level, however, an average of 51.8% sequences of the samples were assigned into the genus-level unclassified genes, and the remaining sequences were generally dominated by Parabacteroides (average relative abundance = 10.6%), Helicobacter (7.6%), Lactobacillus (7.5%), Mucispirillum (6.6%) and diverse genera belonging to other phyla (**Figure 1B**). Taken together, these findings highlight the incomplete coverage of mouse gut microbial genes and taxonomic information in current knowledge. We compared the mouse gut microbial gene catalogue to two comprehensive gene catalogues of the human (9.9 million genes constructed from 1,267 individuals (16)) and rat (5.1 million genes constructed from 98 rats (17)). Only 19.2% genes in the mouse gut microbiotawere also observed in the rat catalogue gut, and only 4.6% genes were shared with the human catalogue (**Figure 1C**). This closer relationship between mouse and rat gut microbiota would be explained by their physiological connections and similar ecological habit, despite that, the low proportion of shared genes between mouse and rat/human gut suggested a unique ecosystem in laboratorial mice.

**Figure 1.**
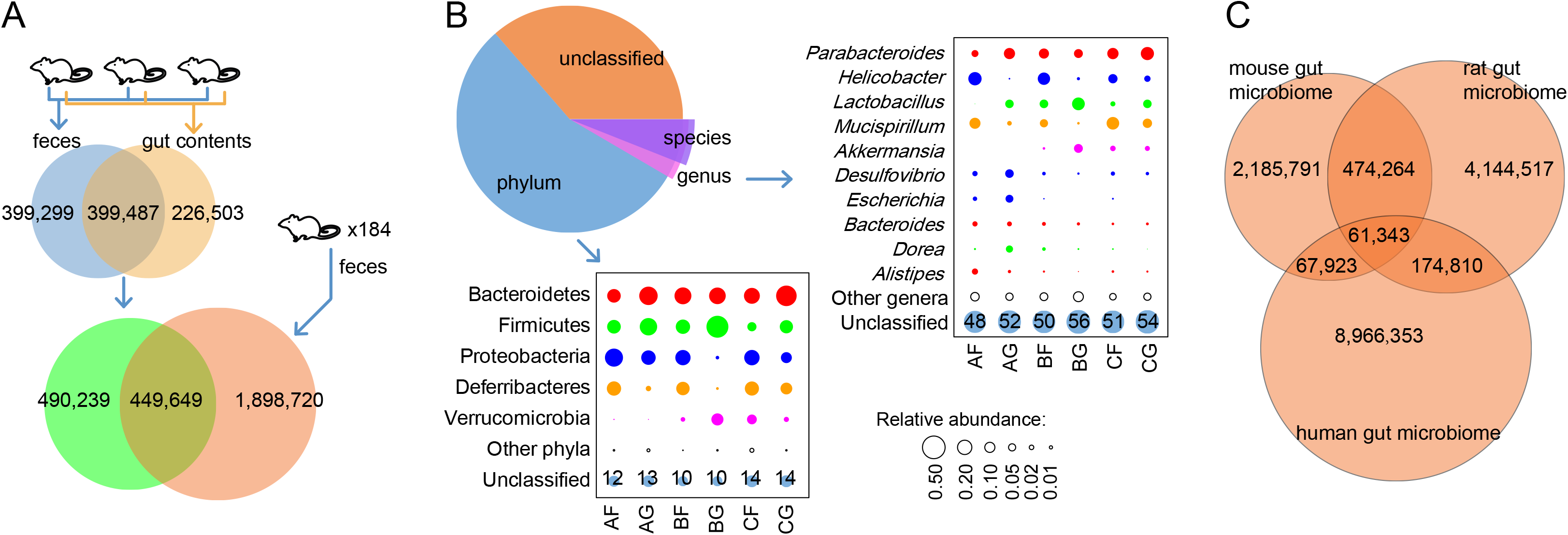
Gene catalogue and microbial community composition of the mouse gut microbiota. **(A)** Comparison of the fecal gene catalogue, the gut content gene catalogue, and the available mouse gut gene catalogue established by 184 mice faeces. **(B)** Microbial community composition of the mouse gut microbiota at the phylum and genus levels. **(C)** Gene sharing relationship of the mouse, rat and human gut microbial gene catalogues.

### Strains resolving of the mouse gut microbiota based on metagenomic-assembled genomes

Recent development of computational techniques allowed us to identify the draft genomes of a portion of uncultivated species in highly diverse metagenomic samples (18, 19). To characterize the microbial strains in mouse intestinal microbiota, we recovered the MAGs for each sample from their assembled contigs (see Materials and Methods for details), which was represented by a total of 236,108 contigs with minimum length of 1.5 kbp (total length of 1.0 Gbp). A total of 112 MAGs were obtained from all samples under strict quality criteria of estimated completeness >70% and contamination <5%. Then, 55 unique MAGs were generated after removing redundancy (**Table S2**), of which 31 MAGs met the standard of high-quality draft genomes (completion >90% and contamination <5%) (**Figure 2A**). These MAGs consisted of 265 scaffolds in average (range from 33 to 760) with an average N50 length of 70 kbp (ranging from 4.2 to 348 kbp), and had an average of sequencing coverage of 96.2X (ranging from 8.1X to 604X). Notably, these 55 MAGs showed distinct genomic similarity with pairwise average nucleotide identity (ANI) ranging from 43% to 95% (average 48%, Figure 2), and captured approximately 50.6% of sequencing reads in all metagenomic-sequenced faecal and gut content samples (**Table S1**), representing a strain-level resolving of the mouse intestinal microbiota.

**Figure 2.**
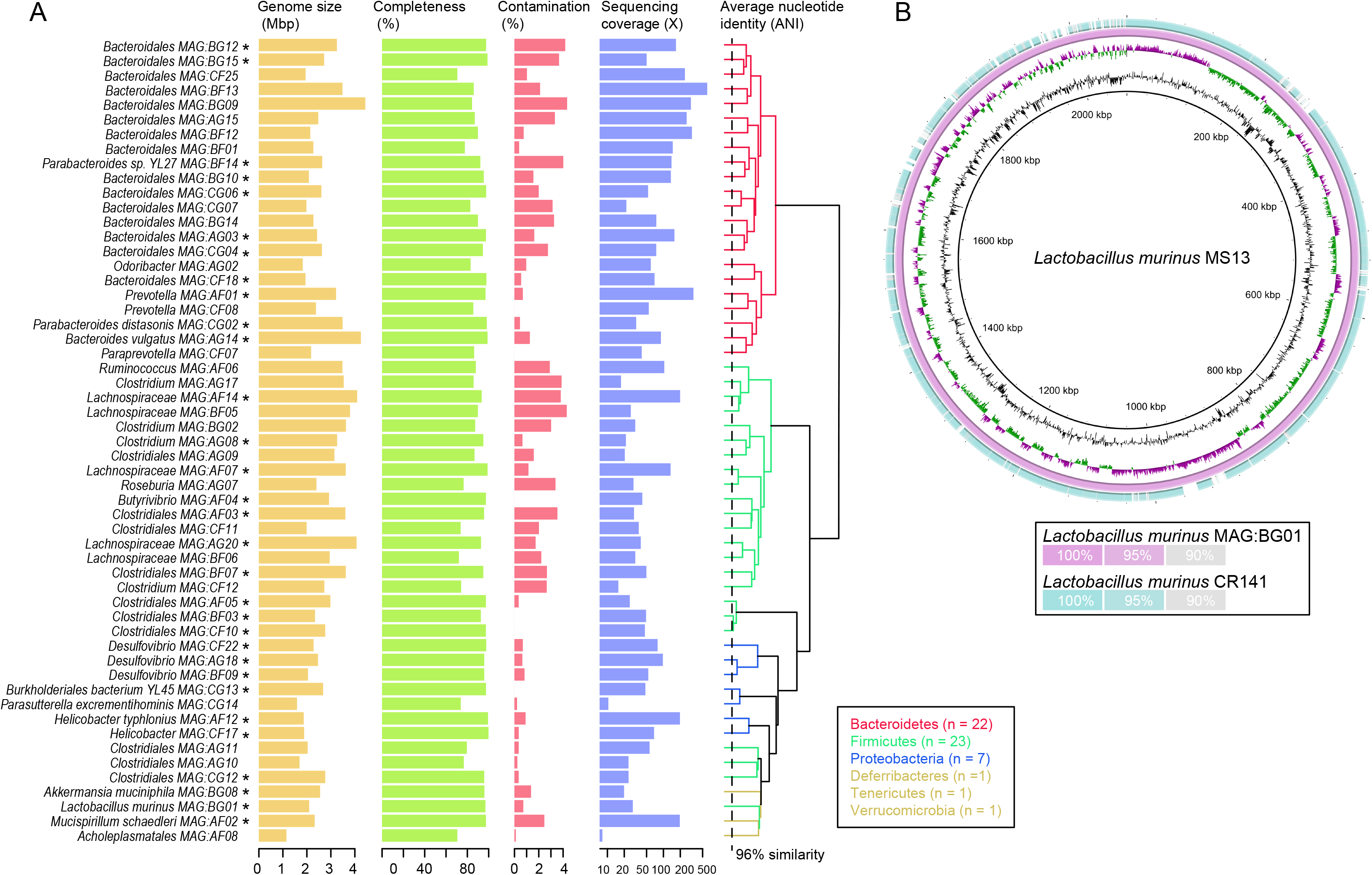
Detailed information and validation of MAGs reconstructed from the mouse gut. **(A)** Summary information of 55 nonredundant MAGs. **(B)** Circular representation of draft genome of a *Lactobacillus murinus* strain isolated from the mouse gut. The inner two circles represent the GC skew and GC content of the *L. murinus* MS13 genome. The outer two circles show the homologous comparison of *L. murinus* MS13 with *L. murinus* MAG:BG01 (reconstructed from the mouse gut) and *L. murinus* CR141 (the closest strain from the NCBI sequenced genomes).

We applied both marker gene-based and whole genome-based approaches to determinate the taxonomic placement of these 55 MAGs. Expectedly, the largest number of strains was from Firmicutes (41.8%, 23/55),Bacteroidetes (40%, 22/55) and Proteobacteria (12.7%, 7/55) (**Figure 2A; Table S2**). And the other strains belonged to relatively low abundance phyla Deferribacteres (n =1), Verrucomicrobia (1) and Tenericutes (1). At the lower taxonomic levels, however, only 9 (16.4%) strains could be robustly assigned to known species, and the remainsing were novel candidate species that were classified into known bacterial genera, families or orders (**Table S2**).

To validate the existence of MAGs and estimate their accuracy, we isolated 13 bacterial colonies via cultivation of a mixed sample from three mice’s faeces and gut contents. Full-length 16S ribosomal RNA gene sequencing was performed on all colonies to enable taxonomic characterization (**Table S3**). The majority of the colonies were members of Gammaproteobacteria and Bacilli, including 4 *Escherichia spp*., 2 *Streptococcus spp*. and 2 *Lactococcus spp*.; these species were of relatively low abundance in the mouse gut microbiota but high cultivability in the common mediums (20). One isolate, *Lactobacillus murinus* MS13, also existed in the MAGs (MAG:BG01). Whole-genome shotgun sequencing analysis of *L. murinus* MS13 revealed an almost identical genome (ANI = 99.99%) compared with MAG:BG01 (**Figure 2B**), confirming that these two genomes were derived from the same strain.

### Distinctive characteristics of mouse’s Bacteroidetes and Firmicutes strains compared to human’s

To investigate the genomic characteristics of the two phyla, we compared our mouse MAGs with 206 strains cultivated from the gastrointestinal tract of healthy adult humans (21) (**Figure 3A; Table S4**). In spite of different gene composition, both mouse and human gut microbiota were dominated by two phyla, Bacteroidetes and Firmicutes (10). The mouse Bacteroidetes strains showed significant reductions of genome size and number of genes, and a significant increase of GC content compared to human Bacteroidetes strains (**Table 1**). In contrast, no significant changes of these parameters were detected between mouse and human Firmicutes strains.

**Figure 3.**
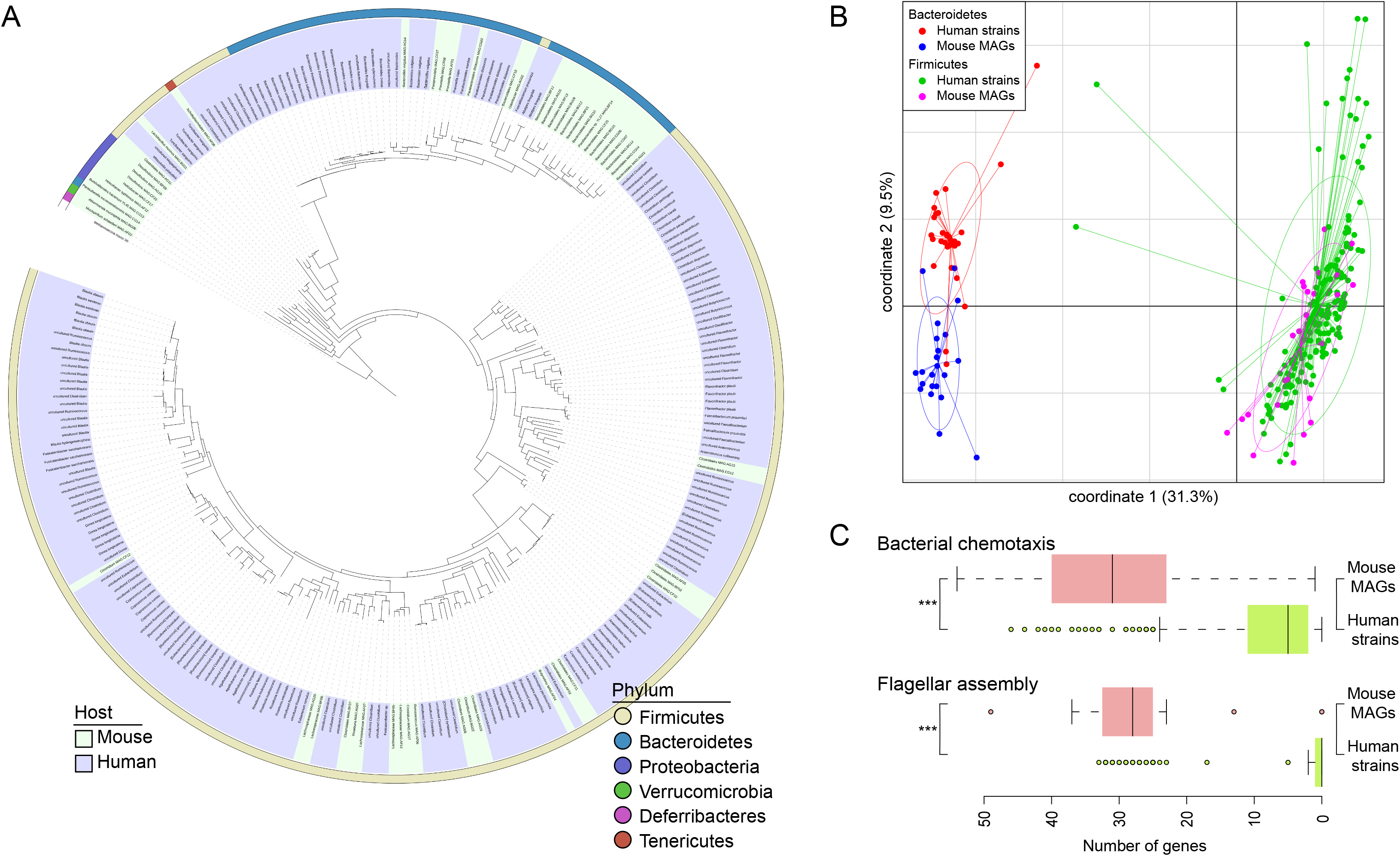
Comparison of Bacteroidetes and Firmicutes strains between mouse and human guts. **(A)** Phylogenetic tree of 55 mouse MAGs and 206 human gut genomes. The outer circle indicates the phylum level taxonomy of the genomes, and the inner circle indicates their host. **(B)** Principle coordination analysis shows the functional difference of Bacteroidetes and Firmicutes strains between mouse and human guts. Isolates on the first and second principal components are plotted by nodes. Lines connect isolates in the same groups, and colored circles cover the isolates near the center of gravity for each group. **(C)** Box-plot shows the difference of flagella assembly and bacterial chemotaxis in the mouse Firmicutes strains compared to human strains.

**Table 1.** Summary of genomic characteristics of Bacteroidetes and Firmicutes strains derived from the human and mouse gut microbiota.

|  | No. of<br>genomes | Genome size<br>(Mbp)* | No. of genes* | % GC<br>content | No. of KEGG<br>orthologs | No. of CAZy<br>proteins |
| --- | --- | --- | --- | --- | --- | --- |
| **Bacteroidetes** |  |  |  |  |  |  |
| Human strains | 31 | 5.1±1.1 | 4,348±908 | 44.8±4.0 | 1558±271 | 357±178 |
| Mouse MAGs | 22 | 2.9±0.8 | 2,637±795 | 50.2±3.3 | 992±176 | 122±49 |
| *P* value |  | 1.8 x 10<sup>-11</sup> | 2.6 x 10<sup>-19</sup> | 1.9 x 10<sup>-6</sup> | 2.1 x 10<sup>-12</sup> | 3.3 x 10<sup>-8</sup> |
| **Firmicutes** |  |  |  |  |  |  |
| Human strains | 175 | 3.7±1.1 | 3,474±1,110 | 43.0±8.1 | 1554±455 | 111±87 |
| Mouse MAGs | 23 | 3.4±0.8 | 3,350±702 | 45.5±6.2 | 1274±224 | 104±46 |
| *P* value |  | 0.122 | 0.464 | 0.094 | 6.7 x 10<sup>-6</sup> | 0.544 |

Note: \* the genome size and number of genes of mouse MAGs were estimated based on their completeness.

To characterize the functional potential of the mouse gut strains, we analyzed the functional profiles of Bacteroidetes and Firmicutes of mouse MAGs and compared them with the corresponding strains of human gut. Consistent with the genomic parameters, the composition of KEGG (Kyoto encyclopedia of genes and genomes (22)) orthologs (KOs) of mouse Bacteroidetes strains demonstrates clear separation from human’s (R^2^ = 0.28, *adonis P* < 0.001; **Figure 3B**), whereas their Firmicutes strains were largely overlapped in KO profiles (R^2^ = 0.039, *adonis P* = 0.01). The mouse Bacteroidetes strains encoded a higher density of enzymes involving genetic information processing compared to human Bacteroidetes strains (average proportion of genes: 20.7% vs. 16.4%, *P* = 1.2 x 10^-11^), and the mouse Firmicutes strains encoded a higher density of enzymes involving cellular processes (average proportion of genes: 12.6% vs. 8.3%, *P* = 8.5 x 10^-10^); whereas strains of the two phyla in mouse gut encoded significantly lower numbers of metabolism-associated enzymes than human strains (*P* = 4.5 x 10^-7^ for Bacteroidetes strains and *P* = 3.8 x 10^-8^ for Firmicutes strains). Such deviation between mouse and human microbial strains was also observed in the low level KEGG pathways, for which 45% (112/248) pathways in Bacteroidetes strains and 23% (63/278) pathways in Firmicutes strains were significantly differed in occurance rates between mouse and human (**Table S5**). As a prominent example, the mouse Firmicutes strains were significantly enriched in pathways of flagella assembly and bacterial chemotaxis compared to human Firmicutes strains (**Figure 3C**).

Consistently, comparison of carbohydrate active enzymes (CAZymes) (23) between mouse MAGs and human strains also revealed significant differences between mouse and human derived Bacteroidetes strains (R^2^ = 0.27, *adonis P* < 0.001), but no significant differences for Firmicutes strains (R^2^ = 0.013, *adonis P* = 0.07). Detailedly, a large proportion (37.5%) of glycoside hydrolases, which were involved hydrolysis and/or rearrangement of glycosidic bonds, were lacked in the mouse Bacteroidetes strains compared to humans’ (**Table S6**).

### Characteristics of other representative strains in the mouse gut microbiota

To study the features of the mouse gut microbes belonging to the other phyla except Bacteroidetes and Firmicutes, we analyzed the genomic characteristics of four MAGs: *Mucispirillum schaedleri* MAG:AF02 from Deferribacteres, *Parasutterella excrementihominis* MAG:CG14 and *Helicobacter typhlonius*MAG:AF12 from Proteobacteria, and *Akkermansia muciniphila* MAG:BG08 from Verrucomicrobia.

*Mucispirillum schaedleri* MAG:AF02 consisted of 47 contigs with total length of 2.32 Mbp (N50 length:166 kbp). This MAG was very similar to the genome of *M. schaedleri* ASF457 with 99.1% ANI (**Figure 4A**), a strain that was isolated from the mucus layer of gastrointestinal tract of laboratory rodents(24).

**Figure 4.**
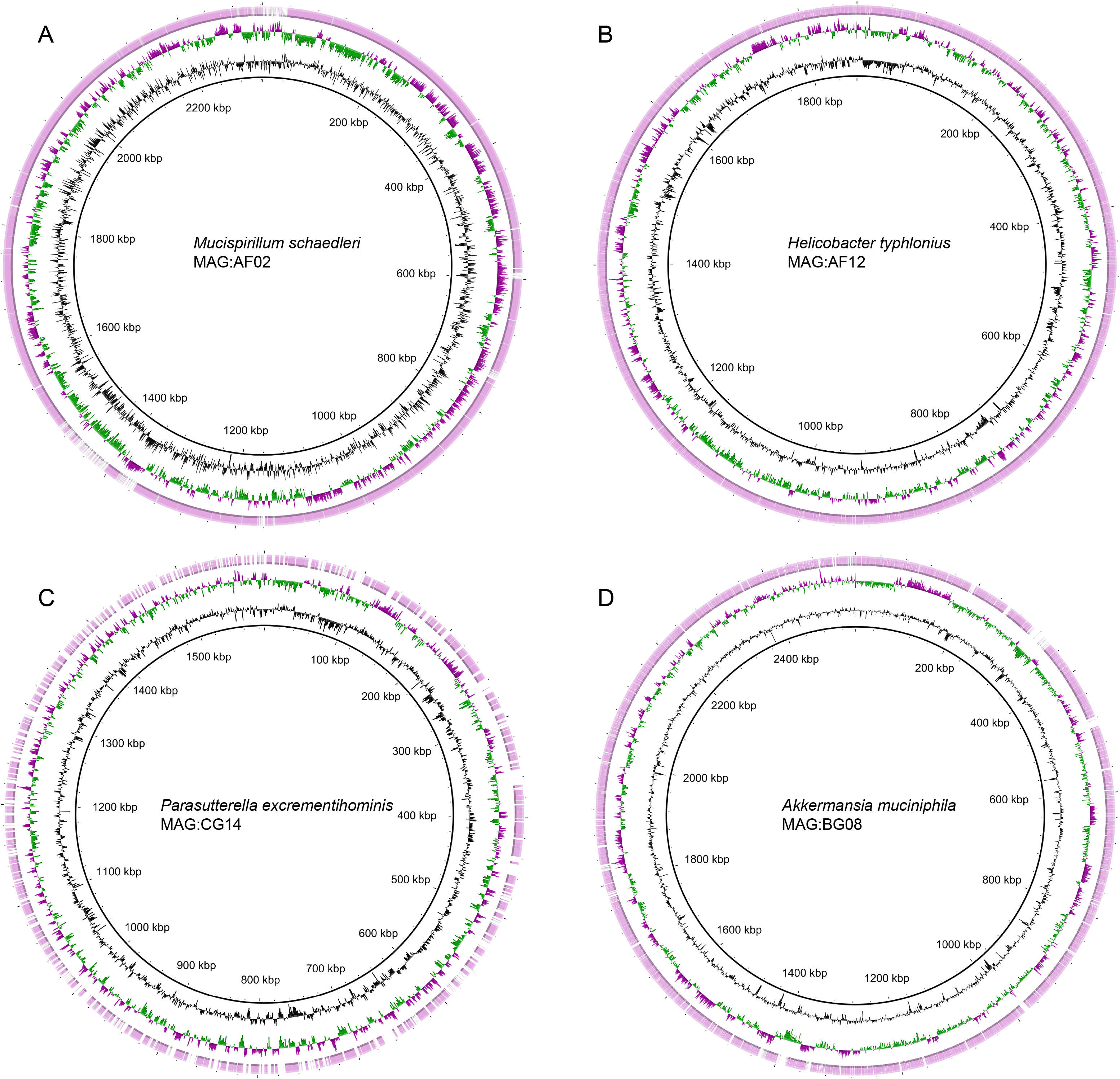
Representative strains in the mouse gut microbiota. Circular representation of draft genome of four representative strains, including *Mucispirillumschaedleri*MAG:AF02 **(A)**, *Parasutterella excrementihominis* MAG:CG14 **(B)**, *Helicobacter typhlonius* MAG:AF12 **(C)**, and *Akkermansia muciniphila* MAG:BG08 **(D)**, in the mouse gut microbiota. The inner two circles represent the GC skew and GC content of the genomes, and the outer circle shows the comparison of the genome with the highest homologous strains from NCBI database.

*Helicobacter typhlonius* MAG:AF12 contained 33 contigs with total length of 1.87 Mbp (N50 length: 237 kbp). The closest genome of MAG:AF12 was *H. typhlonius* mit97-6810 with ANI of 99% (**Figure 4B**).

*Parasutterella excrementihominis* MAG:CG14 contained 397 contigs with total length of 1.59 Mbp. The closest genome of MAG:CG14 was *Sutterella* sp. KGMB03119 with ANI of 96.4% (**Figure 4C**), suggesting that *P. excrementihominis* MAG:CG14 was a potential new sub-species of the genus *Parasutterella*.

*Akkermansia muciniphila* MAG:BG08 contained 216 contigs with total length of 2.55 Mbp. The closest genome of MAG:BG08 was *A. muciniphila* ATCC BAA-835 which was isolated from the human intestinal mucus layer (25), with ANI of 97.7% (**Figure 4D**). This result suggested potentially shared *A.muciniphila* species between mouse and human gut microbiota.

## Discussion

In this study, we compared the mouse gut microbial gene catalogue with that of human and rat, and analyzed the mouse gut microbiota at the strain level based on the metagenome-assembled genome approach. We identified the draft genomes of a portion of uncultivated species from mice faeces and gut content samples. We obtained 55 unique MAGs and 31 of them met the standard of high-quality draft genomes (completion >90% and contamination <5%). Since both mouse and human gut microbiota were dominated by Bacteroidetes and Firmicutes (26, 27), we analyzed these two phyla at the strain level in both, and characterized the potential function of mouse strains through KEGG orthologs and carbohydrate active enzymes. Furthermore, we detected other phyla except Bacteroidetes and Firmicutes, and discovered that some of them have the closest genome with the strains in intestinal mucus layer of laboratory rodents. These outcomes suggested a unique ecosystem in the intestinal tract of laboratorial mice.

According to previous studies, Firmicutes was of the highest abundance in mouse gut microbiota accounting for over 50%, followed by Bacteroidetes, and Deferribacteres and Tenericutes(28). The intestinal bacterial community in mice with acute inflammation may alter with reduced abundance of Firmicutes and Bacteroidetes, especially the clusters Clostridium XIVa and IV (29). Compared with healthy controls, however, Enterobacteriaceae and other clustered groups in the Bacteroidetes may have higher abundance (30–32). Compared to gut microbiota in mouse, the community in human intestinal tract is consistently dominated by Firmicutes and Bacteroidetes. 95% of the Firmicutes sequences are clustered to Clostridia class (10), while most of subspecies of Firmicutes related to butyrate-producing are clustered to clostridial groups IV, XIVa, and XVI (33, 34). In addition, it has been reported that in rat gut microbiota, two phyla Firmicutes and Bacteroidetes have higher abundance, followed by Proteobacteria (17). These outcomes mentioned above are concordant with our findings. Furthermore, we found that 51.8% sequences of our samples belonged to genus-level unclassified genes and the rest of sequences were *Parabacteroides*, *Helicobacter*, *Lactobacillus* and *Mucispirillum*.

Although all specieses we found are assigned to known bacterial genera, families or orders, more than 80% species were classified to novel species based on MAG during our resolving procedure, such as *Acholeplasmatales* MAG:AF08 and *P. excrementihominis* MAG:CG14. It has been reported that uncultivated prokaryotes, such as bacteria, could be obtained by single-amplified genome and MAG approaches (26) and the Genomic Standards Consortium has been completed by minimum information about a MAG, which facilitate robust genomic analyses of bacterial genome (27). However, based on this consequence, the studies on strain-level microbiota in model animals could hardly be found. In addition to analysis by MAG approach, in our study, to confirm the existence of MAGs and to evaluate accuracy, we isolated 13 bacterial colonies via conventional cultivation of a mixed sample from three mice’s faeces and gut contents and all colonies were treated by full-length 16S ribosomal RNA gene sequencing to enable taxonomic characterization. One isolate, *Lactobacillus murinus* MS13, was discovered to have 99.99% average nucleotide identity of its whole genome compared with *L. murinus* MAG:BG01, confirming that these two genomes were derived from the same strain.

The Bacteroidetes and Firmicutes strains of mouse gut showed distinctive functional characteristics compared to that of human gut. Firmicutes and Bacteroidetes are generally Gram-positive and Gramnegative bacteria, respectively. Compared to the single thick and homogeneous layer of Firmicutes, Bacteroidetes has a thinner layered cell wall. The peptidoglycan in the cell walls of Bacteroidetes are less than that in Firmicutes, however, there are lipopolysaccharides, phospholipids, proteins, lipoproteins in the wall of Bacteroidetes and lipids could hardly be found in cell wall of Firmicutes. As the only connection in the elements of cell wall in both two phyla, the chemical composition of peptidoglycan in Bacteroidetes is difficult to be effected(35), in contrast, that of peptidoglycan in Firmicutes could be influenced and variable(36). Thus, this characteristic of peptidoglycan might make Bacteroidetes and Firmicutes dominant in gut microbiota, so as in mouse gut. These pathways of two phyla might be altered when they need to adapt to mouse gut environment.

Apart from Firmicutes and Bacteroidetes, some representative strains are worthy of further study, such as *Akkermansia muciniphila*. *A. muciniphila* is an anaerobic Gram-negative bacteria (37). In previous studies, it has been discovered in the intestines of mice (38). According to our findings, *A. muciniphila* in the mouse gut might have similarities with that in human gut. As an important strain in maintaining healthy gut environment, the alterations of the abundance of *A. muciniphila* may cause many types of diseases, such as type 1 diabetes (39), inflammatory bowel disease(40), autism(41), and cancer(42).

This strain-level study based on MAG approach of mouse gut microbiota might facilitate the development of animal models with more stable physiological performance, benefit clinic diagnosis and therapy and improve the reliability of mouse models. The idea of this research can be applied to many other model organisms, and is of great significance to the selection of organism models and these findings can simulate pathological status of disease more scientifically.

## Materials and Methods

### Experimental models and sample collection

The C57/BL mice were acquired from specific pathogen-free (SPF) Laboratory Animal Center of Dalian Medical University. All mice were housed under a 12:12 light: dark cycle and germ-free conditions in isolators. Animals were fed with autoclaved rodent chow and autoclaved tap water ad libitum. All procedures were performed in accordance with the Guide for the Care and Use of Laboratory Animals under an approved animal study proposal.

To harvest intestinal microbial communities from C57/BL mice, after the mice were sacrificed, the abdomen was sterilized by 75% ethanol and was dissected under sterile conditions. The gut contents and fecal samples were collected for cultivation and DNA extraction.

### Microbial cultivation and identification

Both gut contents and fecal sample of 50 mg were added into 1 ml normal saline and the samples were diluted in BHI liquid medium. The diluents of 20 μl was spread out on solid medium and incubated at 37°C under aerobic and anaerobic conditions for 48 hours, respectively. Then, single colonies were picked up and transferred on fresh solid medium for bacterial purification. After being cultivated under the corresponding conditions for 48 hours, the single colony was inoculated into 8 ml liquid medium and incubated under the same conditions. The culture of 100 μl was used for amplification of 16S rRNA gene by polymerase chain reaction (PCR).

The 16S rRNA gene was amplified by using primers: 7F 5’-AGAGTTTGATYMTGGCTCAG-3’and 1510R 5’-ACGGYTACCTTGTTACGACTT-3’ and the PCR products of 16S rRNA gene were sequenced. Each 16S rRNA sequence was blasted against nucleotide database of NCBI to identify bacterial strains. The identified aerobic strains were preserved in 50% glycerol and anaerobic strains in 50% glycerol and 0.1% cysteine at −80°C freezer.

### DNA extraction, whole-metagenome sequencing, and metagenomic analyses

The microbial DNA of gut content and fecal samples was extracted using Qiagen DNA extraction kit (Qiagen, Germany) according to the manufacturer’s protocols. The DNA concentration and purity were quantified with TBS-380 and NanoDrop2000, respectively. DNA quality was examined with a 1% agarose gel electrophoresis system.

Metagenomic DNA was fragmented to an average size of ~300 bp using Covaris M220 (Gene Company Limited, China). Paired-end libraries were prepared by using a TruSeq DNA sample prep kit (Illumina, USA). Adapters containing the full complement of sequencing primer hybridization sites were ligated to blunt-end fragments. Paired-end sequencing was performed on Illumina HiSeq platform. High-quality reads were extracted based on the FASTQ (43), with default parameters. After then, sequencing reads with >90% similarity to the mouse genomic DNA were removed based on Bowtie2 alignment (44).

A gene catalogue was constructed based on the whole-metagenome sequencing data from the mouse gut samples. High-quality reads were used for *de novo* assembly via MEGAHIT (45), using different k-mer sizes (k = 21, 33, 55, 77). Gene identification was performed for all assembled scaffolds using MetaGeneMark(46). Predicted genes were clustered at the nucleotide level by CD-HIT (47), and genes sharing greater than 90% overlap and greater than 95% identity were treated as redundancies. Taxonomic assignment of the genes was generated by blasting against the NCBI-NT database. When alignments with >70% coverage, genes with >90% and sequence similarity >80% were used for species- and genus-level taxonomical annotation, respectively.

### Metagenome-assembled genome (MAG)

To reconstructed the microbial genome from the metagenomic sequenced samples of mouse gut, we implied the methodology of metagenome-assembled genome that was recently developed by recent studies (18, 48). Briefly, sequenced reads were firstly mapped into the assembled contigs (>2,000bp) with Bowtie2 (44) to generate the mean coverage of contigs. Draft metagenome-assembled genomes were then independently recovered from the contigs of each sample using MetaBAT2 (49) under default parameters, based on the coverage and intrinsic information (e.g. GC content, tetranucleotide frequency) of contigs. The completeness and contamination of the raw MAGs were estimated using CheckM(50), and only MAGs fit the quality criteria of estimated completeness of >70% and contamination of <5% were kept. The pairwise average nucleotide identity (ANI) of MAGs was calculated using FastANI(51), and two MAGs with >95% of ANI were treated as redundancy. Finally, taxonomic assignment of MAGs was performed based on both SpecI (an accurate and universal delineation of prokaryotic species based on marker gene-based algorithm) (52) and whole-genome alignment against the NCBI sequenced bacterial genomes.

### Comparison genomic analyses

Phylogenetic analysis of the genomes was carried out using the maximum-likelihood program RAxML(53) with a GTR model of evolution, and visualized on the iTOL web service (54). Robustness of the phylogenetic tree was estimated by bootstrap analysis in 1,000 replicates. The Kyoto Encyclopedia of Genes and Genomes (KEGG) and carbohydrate active enzymes (CAZymes) databases were used for functional annotation of genomes. For each genome, the protein-coding genes were assigned a KEGG orthologue or CAZyme on the basis of the best-hit gene in the databases.

## Data availability

The raw sequencing data and metagenome-assembled genome sequences reported in this article have been deposited in the NCBI BioProjectPRJNA000000.

## Acknowledgment

This work was supported by National Natural Science Foundation of China (81573469, 81930112, 81902037) and Natural Science Foundation of Liaoning Province, China (20180530086).

## References

1. Richard ML, Sokol H. 2019. The gut mycobiota: insights into analysis, environmental interactions and role in gastrointestinal diseases. Nat Rev Gastroenterol Hepatol 16:331–345.

2. Le Chatelier E, Nielsen T, Qin J, Prifti E, Hildebrand F, Falony G, Almeida M, Arumugam M, Batto JM, Kennedy S, Leonard P, Li J, Burgdorf K, Grarup N, Jorgensen T, Brandslund I, Nielsen HB, Juncker AS, Bertalan M, Levenez F, Pons N, Rasmussen S, Sunagawa S, Tap J, Tims S, Zoetendal EG, Brunak S, Clement K, Dore J, Kleerebezem M, Kristiansen K, Renault P, Sicheritz-Ponten T, de Vos WM, Zucker JD, Raes J, Hansen T, Meta HITc, Bork P, Wang J, Ehrlich SD, Pedersen O. 2013. Richness of human gut microbiome correlates with metabolic markers. Nature 500:541–6.

3. Ley RE, Turnbaugh PJ, Klein S, Gordon JI. 2006. Microbial ecology: human gut microbes associated with obesity. Nature 444:1022–3.

4. Qin J, Li Y, Cai Z, Li S, Zhu J, Zhang F, Liang S, Zhang W, Guan Y, Shen D, Peng Y, Zhang D, Jie Z, Wu W, Qin Y, Xue W, Li J, Han L, Lu D, Wu P, Dai Y, Sun X, Li Z, Tang A, Zhong S, Li X, Chen W, Xu R, Wang M, Feng Q, Gong M, Yu J, Zhang Y, Zhang M, Hansen T, Sanchez G, Raes J, Falony G, Okuda S, Almeida M, LeChatelier E, Renault P, Pons N, Batto JM, Zhang Z, Chen H, Yang R, Zheng W, Li S, Yang H, et al. 2012. A metagenome-wide association study of gut microbiota in type 2 diabetes. Nature 490:55–60.

5. Vaahtovuo J, Munukka E, Korkeamaki M, Luukkainen R, Toivanen P. 2008. Fecal microbiota in early rheumatoid arthritis. J Rheumatol 35:1500–5.

6. Russell SL, Gold MJ, Hartmann M, Willing BP, Thorson L, Wlodarska M, Gill N, Blanchet MR, Mohn WW, McNagny KM, Finlay BB. 2012. Early life antibiotic-driven changes in microbiota enhance susceptibility to allergic asthma. EMBO Rep 13:440–7.

7. Tlaskalova-Hogenova H, Stepankova R, Kozakova H, Hudcovic T, Vannucci L, Tuckova L, Rossmann P, Hrncir T, Kverka M, Zakostelska Z, Klimesova K, Pribylova J, Bartova J, Sanchez D, Fundova P, Borovska D, Srutkova D, Zidek Z, Schwarzer M, Drastich P, Funda DP. 2011. The role of gut microbiota (commensal bacteria) and the mucosal barrier in the pathogenesis of inflammatory and autoimmune diseases and cancer: contribution of germ-free and gnotobiotic animal models of human diseases. Cell Mol Immunol 8:110–20.

8. Adak A, Khan MR. 2019. An insight into gut microbiota and its functionalities. Cell Mol Life Sci 76:473–493.

9. Nguyen TL, Vieira-Silva S, Liston A, Raes J. 2015. How informative is the mouse for human gut microbiota research? Dis Model Mech 8:1–16.

10. Eckburg PB, Bik EM, Bernstein CN, Purdom E, Dethlefsen L, Sargent M, Gill SR, Nelson KE, Relman DA. 2005. Diversity of the human intestinal microbial flora. Science 308:1635–8.

11. Ley RE, Backhed F, Turnbaugh P, Lozupone CA, Knight RD, Gordon JI. 2005. Obesity alters gut microbial ecology. Proc Natl Acad Sci U S A 102:11070–5.

12. Dumont BL. 2019. Significant Strain Variation in the Mutation Spectra of Inbred Laboratory Mice. Mol Biol Evol 36:865–874.

13. Gu S, Chen D, Zhang JN, Lv X, Wang K, Duan LP, Nie Y, Wu XL. 2013. Bacterial community mapping of the mouse gastrointestinal tract. PLoS One 8:e74957.

14. Xiao L, Feng Q, Liang S, Sonne SB, Xia Z, Qiu X, Li X, Long H, Zhang J, Zhang D, Liu C, Fang Z, Chou J, Glanville J, Hao Q, Kotowska D, Colding C, Licht TR, Wu D, Yu J, Sung JJ, Liang Q, Li J, Jia H, Lan Z, Tremaroli V, Dworzynski P, Nielsen HB, Backhed F, Dore J, Le Chatelier E, Ehrlich SD, Lin JC, Arumugam M, Wang J, Madsen L, Kristiansen K. 2015. A catalog of the mouse gut metagenome. Nat Biotechnol 33:1103–8.

15. Hildebrand F, Nguyen TL, Brinkman B, Yunta RG, Cauwe B, Vandenabeele P, Liston A, Raes J. 2013. Inflammation-associated enterotypes, host genotype, cage and inter-individual effects drive gut microbiota variation in common laboratory mice. Genome Biol 14:R4.

16. Li J, Jia H, Cai X, Zhong H, Feng Q, Sunagawa S, Arumugam M, Kultima JR, Prifti E, Nielsen T, Juncker AS, Manichanh C, Chen B, Zhang W, Levenez F, Wang J, Xu X, Xiao L, Liang S, Zhang D, Zhang Z, Chen W, Zhao H, Al-Aama JY, Edris S, Yang H, Wang J, Hansen T, Nielsen HB, Brunak S, Kristiansen K, Guarner F, Pedersen O, Dore J, Ehrlich SD, Meta HITC, Bork P, Wang J, Meta HITC. 2014. An integrated catalog of reference genes in the human gut microbiome. Nat Biotechnol 32:834–41.

17. Pan H, Guo R, Zhu J, Wang Q, Ju Y, Xie Y, Zheng Y, Wang Z, Li T, Liu Z, Lu L, Li F, Tong B, Xiao L, Xu X, Li R, Yuan Z, Yang H, Wang J, Kristiansen K, Jia H, Liu L. 2018. A gene catalogue of the Sprague-Dawley rat gut metagenome. Gigascience 7.

18. Pasolli E, Asnicar F, Manara S, Zolfo M, Karcher N, Armanini F, Beghini F, Manghi P, Tett A, Ghen-si P, Collado MC, Rice BL, DuLong C, Morgan XC, Golden CD, Quince C, Huttenhower C, Segata N. 2019. Extensive Unexplored Human Microbiome Diversity Revealed by Over 150,000 Genomes from Metagenomes Spanning Age, Geography, and Lifestyle. Cell 176:649–662 e20.

19. Nayfach S, Shi ZJ, Seshadri R, Pollard KS, Kyrpides NC. 2019. New insights from uncultivated genomes of the global human gut microbiome. Nature 568:505–510.

20. Lagkouvardos I, Overmann J, Clavel T. 2017. Cultured microbes represent a substantial fraction of the human and mouse gut microbiota. Gut Microbes 8:493–503.

21. Browne HP, Forster SC, Anonye BO, Kumar N, Neville BA, Stares MD, Goulding D, Lawley TD. 2016. Culturing of ‘unculturable’ human microbiota reveals novel taxa and extensive sporulation. Nature 533:543–546.

22. Kanehisa M, Furumichi M, Tanabe M, Sato Y, Morishima K. 2017. KEGG: new perspectives on genomes, pathways, diseases and drugs. Nucleic Acids Res 45:D353–D361.

23. Lombard V, Golaconda Ramulu H, Drula E, Coutinho PM, Henrissat B. 2014. The carbohydrateactive enzymes database (CAZy) in 2013. Nucleic Acids Res 42:D490–5.

24. Robertson BR, O’Rourke JL, Neilan BA, Vandamme P, On SL, Fox JG, Lee A. 2005. Mucispirillum schaedleri gen. nov., sp. nov., a spiral-shaped bacterium colonizing the mucus layer of the gastrointestinal tract of laboratory rodents. Int J Syst Evol Microbiol 55:1199–204.

25. Derrien M, Vaughan EE, Plugge CM, de Vos WM. 2004. Akkermansia muciniphila gen. nov., sp. nov., a human intestinal mucin-degrading bacterium. Int J Syst Evol Microbiol 54:1469–76.

26. Alneberg J, Karlsson CMG, Divne AM, Bergin C, Homa F, Lindh MV, Hugerth LW, Ettema TJG, Bertilsson S, Andersson AF, Pinhassi J. 2018. Genomes from uncultivated prokaryotes: a comparison of metagenome-assembled and single-amplified genomes. Microbiome 6:173.

27. Bowers RM, Kyrpides NC, Stepanauskas R, Harmon-Smith M, Doud D, Reddy TBK, Schulz F, Jarett J, Rivers AR, Eloe-Fadrosh EA, Tringe SG, Ivanova NN, Copeland A, Clum A, Becraft ED, Malmstrom RR, Birren B, Podar M, Bork P, Weinstock GM, Garrity GM, Dodsworth JA, Yooseph S, Sutton G, Glockner FO, Gilbert JA, Nelson WC, Hallam SJ, Jungbluth SP, Ettema TJG, Tighe S, Konstantinidis KT, Liu WT, Baker BJ, Rattei T, Eisen JA, Hedlund B, McMahon KD, Fierer N, Knight R, Finn R, Cochrane G, Karsch-Mizrachi I, Tyson GW, Rinke C, Genome Standards C, Lapidus A, Meyer F, Yilmaz P, Parks DH, et al. 2017. Minimum information about a single amplified genome (MISAG) and a metagenome-assembled genome (MIMAG) of bacteria and archaea. Nat Biotechnol 35:725–731.

28. Rosshart SP, Vassallo BG, Angeletti D, Hutchinson DS, Morgan AP, Takeda K, Hickman HD, McCulloch JA, Badger JH, Ajami NJ, Trinchieri G, Pardo-Manuel de Villena F, Yewdell JW, Rehermann B. 2017. Wild Mouse Gut Microbiota Promotes Host Fitness and Improves Disease Resistance. Cell 171:1015–1028 e13.

29. Robertson SJ, Goethel A, Girardin SE, Philpott DJ. 2018. Innate Immune Influences on the Gut Microbiome: Lessons from Mouse Models. Trends Immunol 39:992–1004.

30. Schwab C, Berry D, Rauch I, Rennisch I, Ramesmayer J, Hainzl E, Heider S, Decker T, Kenner L, Muller M, Strobl B, Wagner M, Schleper C, Loy A, Urich T. 2014. Longitudinal study of murine microbiota activity and interactions with the host during acute inflammation and recovery. ISME J 8:1101–14.

31. Berry D, Schwab C, Milinovich G, Reichert J, Ben Mahfoudh K, Decker T, Engel M, Hai B, Hainzl E, Heider S, Kenner L, Muller M, Rauch I, Strobl B, Wagner M, Schleper C, Urich T, Loy A. 2012. Phylotype-level 16S rRNA analysis reveals new bacterial indicators of health state in acute murine colitis. ISME J 6:2091–106.

32. Robertson SJ, Geddes K, Maisonneuve C, Streutker CJ, Philpott DJ. 2016. Resilience of the intestinal microbiota following pathogenic bacterial infection is independent of innate immunity mediated by NOD1 or NOD2. Microbes Infect 18:460–71.

33. Barcenilla A, Pryde SE, Martin JC, Duncan SH, Stewart CS, Henderson C, Flint HJ. 2000. Phylogenetic relationships of butyrate-producing bacteria from the human gut. Appl Environ Microbiol 66:1654–61.

34. Pryde SE, Duncan SH, Hold GL, Stewart CS, Flint HJ. 2002. The microbiology of butyrate formation in the human colon. FEMS Microbiol Lett 217:133–9.

35. Schleifer KH, Kandler O. 1972. Peptidoglycan types of bacterial cell walls and their taxonomic implications. Bacteriol Rev 36:407–77.

36. Schleifer KH. 2009. Classification of Bacteria and Archaea: past, present and future. Syst Appl Microbiol 32:533–42.

37. Xing J, Li X, Sun Y, Zhao J, Miao S, Xiong Q, Zhang Y, Zhang G. 2019. Comparative genomic and functional analysis of Akkermansia muciniphila and closely related species. Genes Genomics 41:1253–1264.

38. Presley LL, Wei B, Braun J, Borneman J. 2010. Bacteria associated with immunoregulatory cells in mice. Appl Environ Microbiol 76:936–41.

39. Hansen CH, Krych L, Nielsen DS, Vogensen FK, Hansen LH, Sorensen SJ, Buschard K, Hansen AK. 2012. Early life treatment with vancomycin propagates Akkermansia muciniphila and reduces diabetes incidence in the NOD mouse. Diabetologia 55:2285–94.

40. Png CW, Linden SK, Gilshenan KS, Zoetendal EG, McSweeney CS, Sly LI, McGuckin MA, Florin TH. 2010. Mucolytic bacteria with increased prevalence in IBD mucosa augment in vitro utilization of mucin by other bacteria. Am J Gastroenterol 105:2420–8.

41. Wang L, Christophersen CT, Sorich MJ, Gerber JP, Angley MT, Conlon MA. 2011. Low relative abundances of the mucolytic bacterium Akkermansia muciniphila and Bifidobacterium spp. in feces of children with autism. Appl Environ Microbiol 77:6718–21.

42. Weir TL, Manter DK, Sheflin AM, Barnett BA, Heuberger AL, Ryan EP. 2013. Stool microbiome and metabolome differences between colorectal cancer patients and healthy adults. PLoS One 8:e70803.

43. Chen S, Zhou Y, Chen Y, Gu J. 2018. fastp: an ultra-fast all-in-one FASTQ preprocessor. Bioinformatics 34:i884–i890.

44. Langmead B, Salzberg SL. 2012. Fast gapped-read alignment with Bowtie 2. Nat Methods 9:357–9.

45. Li D, Liu CM, Luo R, Sadakane K, Lam TW. 2015. MEGAHIT: an ultra-fast single-node solution for large and complex metagenomics assembly via succinct de Bruijn graph. Bioinformatics 31:1674–6.

46. Zhu W, Lomsadze A, Borodovsky M. 2010. Ab initio gene identification in metagenomic sequences. Nucleic Acids Res 38:e132.

47. Li W, Godzik A. 2006. Cd-hit: a fast program for clustering and comparing large sets of protein or nucleotide sequences. Bioinformatics 22:1658–9.

48. Parks DH, Rinke C, Chuvochina M, Chaumeil PA, Woodcroft BJ, Evans PN, Hugenholtz P, Tyson GW. 2017. Recovery of nearly 8,000 metagenome-assembled genomes substantially expands the tree of life. Nat Microbiol 2:1533–1542.

49. Kang DD, Li F, Kirton E, Thomas A, Egan R, An H, Wang Z. 2019. MetaBAT 2: an adaptive binning algorithm for robust and efficient genome reconstruction from metagenome assemblies. PeerJ 7:e7359.

50. Parks DH, Imelfort M, Skennerton CT, Hugenholtz P, Tyson GW. 2015. CheckM: assessing the quality of microbial genomes recovered from isolates, single cells, and metagenomes. Genome Res 25:1043–55.

51. Jain C, Rodriguez RL, Phillippy AM, Konstantinidis KT, Aluru S. 2018. High throughput ANI analysis of 90K prokaryotic genomes reveals clear species boundaries. Nat Commun 9:5114.

52. Mende DR, Sunagawa S, Zeller G, Bork P. 2013. Accurate and universal delineation of prokaryotic species. Nat Methods 10:881–4.

53. Stamatakis A. 2014. RAxML version 8: a tool for phylogenetic analysis and post-analysis of large phylogenies. Bioinformatics 30:1312–3.

54. Letunic I, Bork P. 2016. Interactive tree of life (iTOL) v3: an online tool for the display and annotation of phylogenetic and other trees. Nucleic Acids Res 44:W242–5.

